# SUMO protease FUG1, histone reader AL3 and the PRC1 Complex are integral to repeat-expansion induced epigenetic silencing in *Arabidopsis thaliana*

**DOI:** 10.1101/2023.01.13.523841

**Authors:** Sridevi Sureshkumar, Champa Bandaranayake, Junqing Lu, Craig I Dent, Chhaya Atri, Harrison M York, Prashanth Tamizhselvan, Nawar Shamaya, Giulia Folini, Prakash Kumar Bhagat, Benjamin G Bergey, Avilash Singh Yadav, Subhasree Kumar, Oliver Grummisch, Prince Saini, Ram K Yadav, Senthil Arumugam, Emanuel Rosonina, Ari Sadanandom, Hongtao Liu, Sureshkumar Balasubramanian

## Abstract

Epigenetic gene silencing induced by expanded repeats can cause diverse phenotypes ranging from severe growth defects in plants to genetic diseases such as Friedreich’s ataxia in humans^1^. The molecular mechanisms underlying repeat expansion-induced epigenetic silencing remain largely unknown^2,3^. Using a plant model, we have previously shown that expanded repeats can induce smallRNAs which in turn can lead to epigenetic silencing through the RNA-dependent DNA methylation pathway^4,5^. Here, using a genetic suppressor screen, we confirm a key role for the RdDM pathway and identify novel components required for epigenetic silencing caused by expanded repeats. We show that FOURTH ULP LIKE GENE CLASS 1 (FUG1) – a SUMO protease, ALFIN-LIKE 3 – a histone reader and LIKE HETEROCHROMATIN 1 (LHP1) - a component of the PRC1 complex are required for repeat expansion-induced epigenetic silencing. Loss of any of these components suppress repeat expansion-associated phenotypes. SUMO protease FUG1 physically interacts with AL3 and perturbing its potential SUMOylation site disrupts its nuclear localisation. AL3 physically interacts with LHP1 of the PRC1 complex and the FUG1-AL3-LHP1 module is essential to confer repeat expansion-associated epigenetic silencing. Our findings highlight the importance post-translational modifiers and histone readers in epigenetic silencing caused by repeat expansions.

---

Variation in the length of short tandem repeats is associated with diverse phenotypic changes across organisms^6-11^. In humans, extreme length variants exemplified by repeat expansions cause several diseases such as Huntington’s disease and Friedreich’s ataxia^8,12^. Recent explosion in genomic data has resulted in the discovery of novel repeat expansion-associated phenotypes in multiple organisms and new diseases including cancer^13-16^. Gene silencing associated with repeat expansions is seen in diseases like Friedreich’s ataxia, but how repeat expansions lead to epigenetic silencing remains largely unknown^1,2,17^.

The Bur-0 accession of Arabidopsis carries a GAA/TTC repeat expansion in the 3^rd^ intron of the *ISOPROPYLMALATE ISOMERASE LARGE SUBUNIT 1 (IIL1)* gene associated with the epigenetic silencing at the *IIL1* locus^5^. Triplet expansion-induced reduction in *IIL1* expression leads to a temperature-sensitive growth defect referred to as the *irregularly impaired leaves (iil)* phenotype (narrower and twisted leaves) when grown at 27ºC^5^. We have previously shown that the GAA/TTC repeat expansion at this locus leads to accumulation of 24nt siRNAs that map to *IIL1*, and through the RdDM pathway leads to epigenetic silencing^4^.

To decipher molecular pathways that mediate repeat expansion-associated gene silencing, we carried out a new large-scale EMS mutagenesis screen in the Bur-0 (*iil* mutant) background (Fig. 1a). We quantified *IIL1* expression to identify suppressors that act at the level of gene expression and identified a total of nine suppressors with elevated *IIL1* levels (Fig. 1b). F_1_ progeny of all nine genetic suppressors in a cross with Bur-0 revealed that they were recessive in nature. We sequenced the mutants and used a combination of SHOREMAP^18^ and positional cloning to identify the potential underlying mutations in these nine genetic suppressors of *iil*. For four mutants, we sequenced a pool of 500 segregating plants from an F_2_ population and mapped potential loss-of-function alleles at *At3g57080* (*RPB5B*, an RNA *Pol V* subunit in *44-2*), *At2g40030* (*NRPD1B*, an RNA *Pol V* subunit in *49-9*), *At2g27040* (AGO4 in *57-3*) and a mis-sense mutation at *At3g23780* (*NRPD2A*, an *RNA Pol IV* subunit in *61-7*) (Fig 1c and Fig S1). We have previously shown that knocking down *At2g40030, At3g23780 and At2g27040* in the Bur-0 background suppresses the *iil* phenotype and therefore, we conclude that the observed mutations are causal. Since SHOREMAP analysis and our earlier findings^4^ confirmed the importance of the RdDM pathway for repeat expansion-induced gene silencing, we analysed the rest of the mutants by direct sequencing for mutations in RdDM pathway genes. These analyses revealed that four other suppressors harboured mutations in *At4g11130* (*RDR2* in 71-9) or in *At2g27040* (*AGO4* in *72-6, 81-2* and *108-8*) (Supplementary Table ST1). In summary of the nine genetic suppressors, 8 were mutants in RdDM pathway genes. We conclude that RdDM is the major pathway by which expanded repeats cause epigenetic silencing in *Arabidopsis thaliana*.

**Figure 1.**
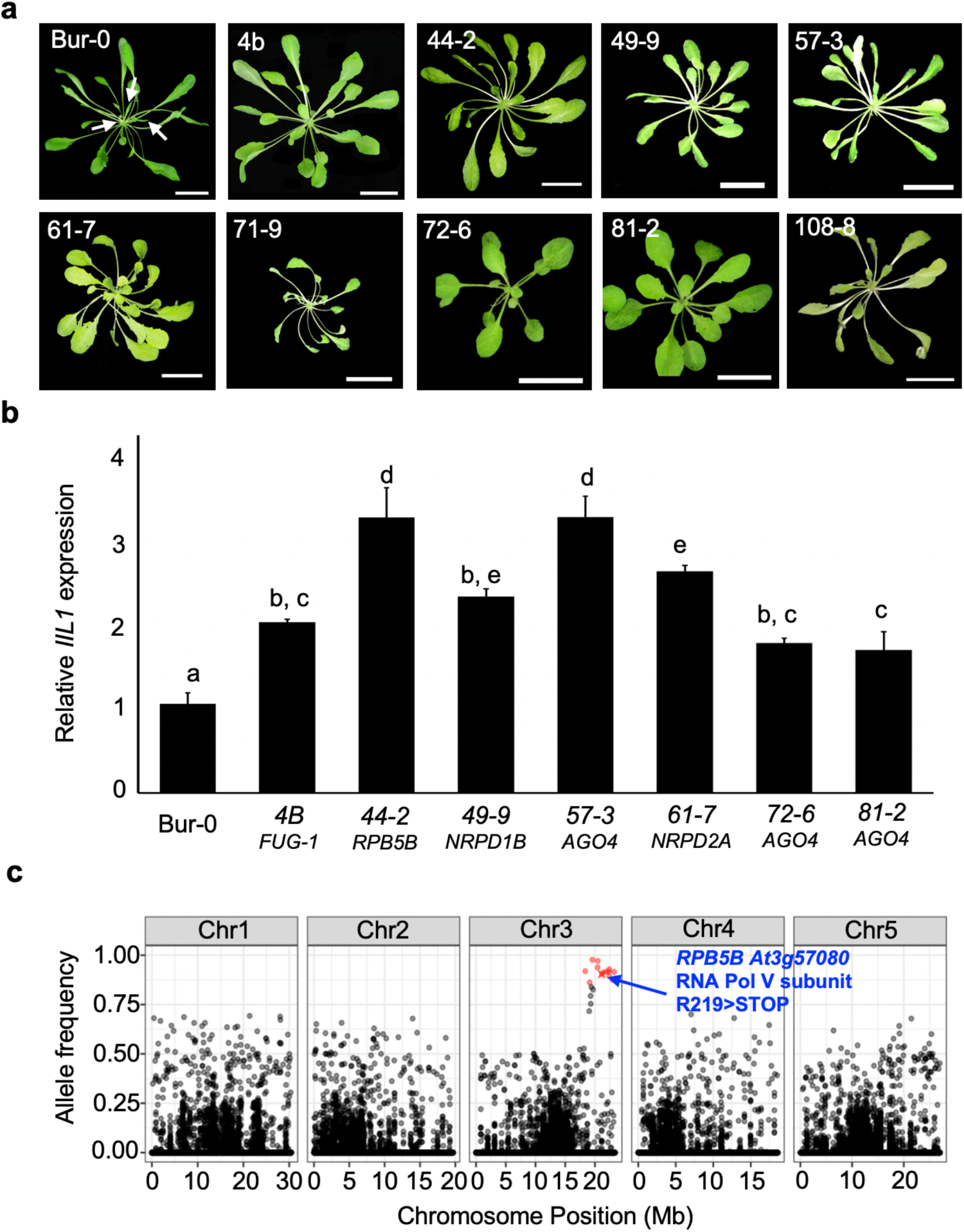
A genetic suppressor screen identifies RdDM to be the major pathway for repeat expansion-induced epigenetic silencing. a) Phenotypes of the isolated suppressors compared to Bur-0. The “*irregularly impaired leaves”* are marked by white arrows in the Bur-0 wild type. Scale bars represent 2 cm. b) Relative *IIL1* expression levels in genetic suppressors identified through the genetic screen. The numbers represent the original screen identifiers and the corresponding genes identified after cloning are shown below. Average expression levels based on at least two biological replicates for each line are shown. p-values are based on Tukey’s post hoc test, and lines that are not harbouring the same letters are significantly different from each other (p<0.05). Error bars represent SEM. c) An example of SHOREMAP analysis with 44-2 identifies a mutation in Pol V. High frequency alleles (>0.85) are coloured red and red crosses show the putative causal alleles.

One mutant (*4b*) did not carry mutations in the candidates for the RdDM pathway (Fig 1a and Fig S2). Sequence analysis failed to discover any major-effect mutations within a 250Kb region surrounding the *IIL1* locus, which indicated that *4b* is a second-site suppressor. We detected 34 potential loss of function mutations in *4b* but none of them were in obvious candidate genes (Supplementary Table ST2). We crossed the *4b* mutant with Pf-0 (*IIL* wild type), since the original repeat-expansion-associated *iil* phenotype in the Bur-0 background segregated as a monogenic trait in this cross^5^. We then selected lines that are homozygous for *IIL1* and still harbouring the repeat expansions and used the progeny of these lines for mapping *4b*. By linkage mapping using a total of more than 750 mutant plants, we mapped *4b* to a 90 kb region on chromosome 3, containing 20 protein coding genes (Fig 2a). We identified only a single EMS-type SNP within this 90 kb that was in the first exon of *At3g48480* at position 17958141 (corresponding to TAIR10 in Col-0), which changes Gln into a stop codon (Q152*) making it a potential candidate gene (Fig 2a). *At3g48480* encodes a protein referred to as FOURTH UBIQUITIN LIKE GENE CLASS 1 (FUG1)^19^. To test whether *FUG1* is *4b*, we designed two independent artificial microRNAs (amiRNAs, *35S::amiR-FUG1*) and generated knockdown lines in the Bur-0 (*iil* mutant) background (Fig 2b). 190 of the 200 independent *35S::amiR-FUG1* lines in the Bur-0 background displayed suppression of the *iil* phenotype (Fig 2b), confirming that the loss of function of *FUG1* is sufficient to suppress the *iil* phenotype. Conversely, overexpression of *FUG1* (*35S::FUG1*) in the *4b* background restored the *iil* phenotype (Fig 2b). Henceforth the *4b* will be referred to as the *fug1* mutant.

**Figure 2.**
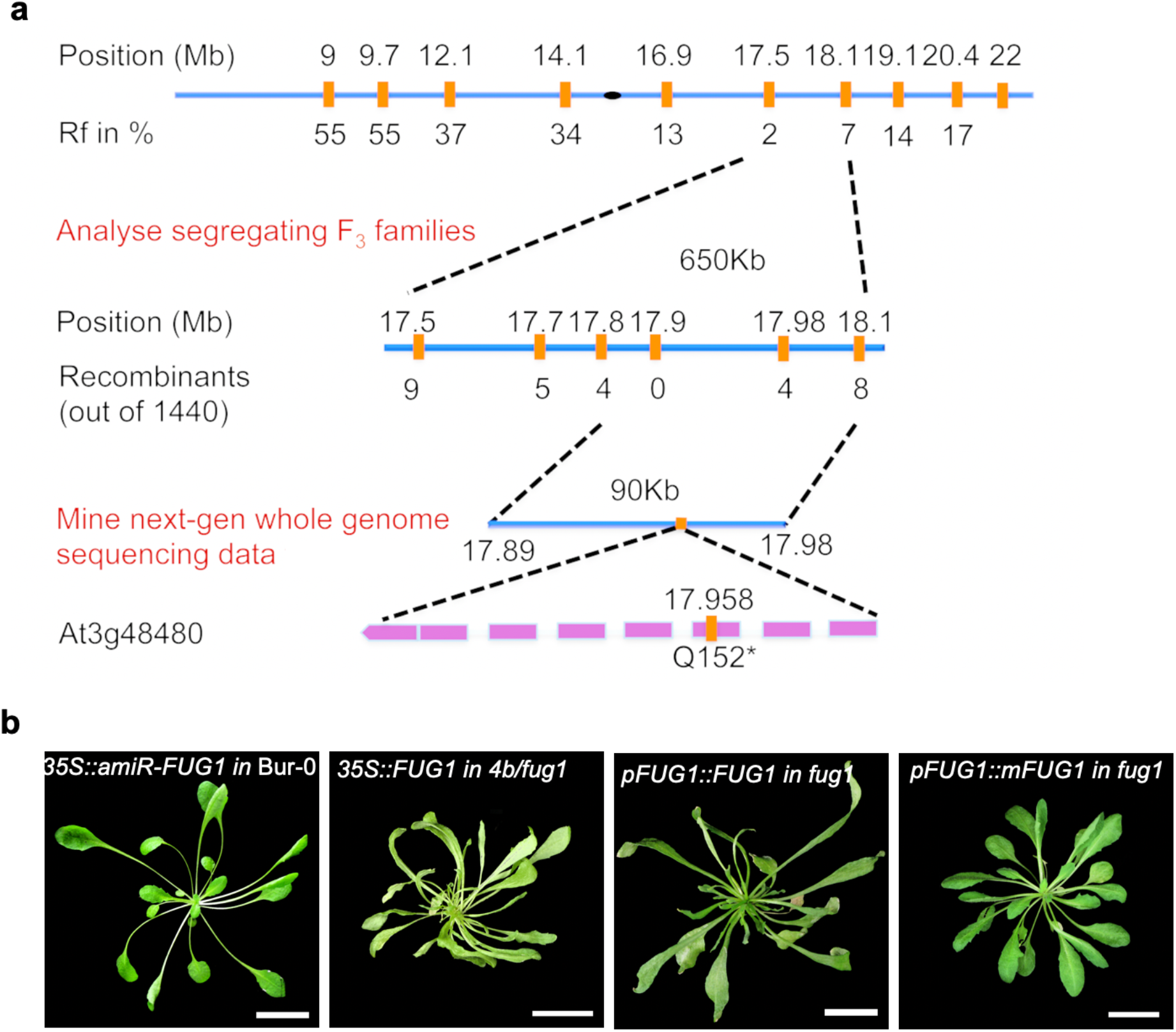
Positional cloning of the *4b/fug1* identifies a SUMO protease. a) Positional cloning of the *4b/fug1* mutant. Analysis at F_2_ and F_3_ are shown in two stages and subsequently the 90Kb interval was analysed through sequencing. b) At3g48480 is FUG1 and its protease domain is essential for its function. Phenotypes of *35S::amiR-FUG1* in Bur-0, *35S::FUG1 in fug1, pFUG1::FUG1 in fug1* as well as *pFUG1::mFUG1 in fug1* are shown. Scale bars represent 2 cm.

FUG1 belongs to a highly conserved superfamily of cysteine-type proteases and its related members in Arabidopsis are potential Small Ubiquitin-like Modifier (SUMO) proteases involved in deSUMOylation^19,20^. Among the SUMO proteases in Arabidopsis FUG1 remains an uncharacterized protein^20^. Mutants for the rice orthologue of FUG1 show an increase in SUMOylation levels^21^, which suggests that FUG1 could function as a deSUMOylase. We used the heterologous yeast system to test whether the protease domain of the Arabidopsis FUG1 is functional by swapping it with the same of yeast Ubiquitin like-Specific-protease 2 (ULP2). Yeast strains with mutations in *ULP2 (ulp2Δ)* or *ULP1 (ulp1-I615N)* grow normally at 30ºC but are severely compromised in growth at 37ºC ^22^. While *FUG1* did not complement this growth defect (Fig S3a-b), a partial recovery was seen in yeast cells harbouring hybrid constructs in which the protease domain of ULP2 was swapped with that of FUG1 in liquid cultures (Fig S3c-f). We observed several high-molecular weight proteins suggestive of poly-SUMOylation upon mutations in *ULP2*, which was reduced in cells harbouring the hybrid construct (Fig S3g). These findings suggested that FUG1 has a functional protease domain increasing its likelihood of being a deSUMOylase.

To test whether the protease function of FUG1 is required for the observed *iil* phenotype, we disrupted the catalytic triad in the FUG1 protein (C246S) and generated transgenic plants expressing either wild type *FUG1 (pFUG1:FUG1)* or mutated *FUG1 (pFUG1::mFUG1)* under its native promoter and transformed them into the *fug1* mutant background. *pFUG1::FUG1* plants in the *fug1* background displayed the *iil* phenotype (restoring to Bur-0 situation) at 27ºC demonstrating that the construct is functional (Fig 2b). Conversely, *pFUG1::mFUG1* plants failed to complement the *fug1* mutant (normal looking plant) confirming the requirement of the protease domain for FUG1 function (Fig 2b). We conclude that FUG1 encodes a SUMO protease that is required to confer repeat-expansion associated gene silencing.

FUG1 was localized in the nucleus of *35S::GFP-FUG1* plants (Fig S4). To identify potential FUG1 interactors, we performed a yeast two-hybrid screen and recovered a plant homeodomain (PHD) finger containing nuclear protein ALFIN LIKE 3 (AL3)^23^. PHD finger containing proteins are known to be histone H3K4me2/3 readers^24,25^. FUG1-AL3 interaction was confirmed in yeast cells (Fig 3a). Split-GFP assays with co-transfection on tobacco leaf epidermal cells confirmed the *in planta* nuclear interaction of FUG1 and AL3 (Fig 3b).

**Figure 3.**
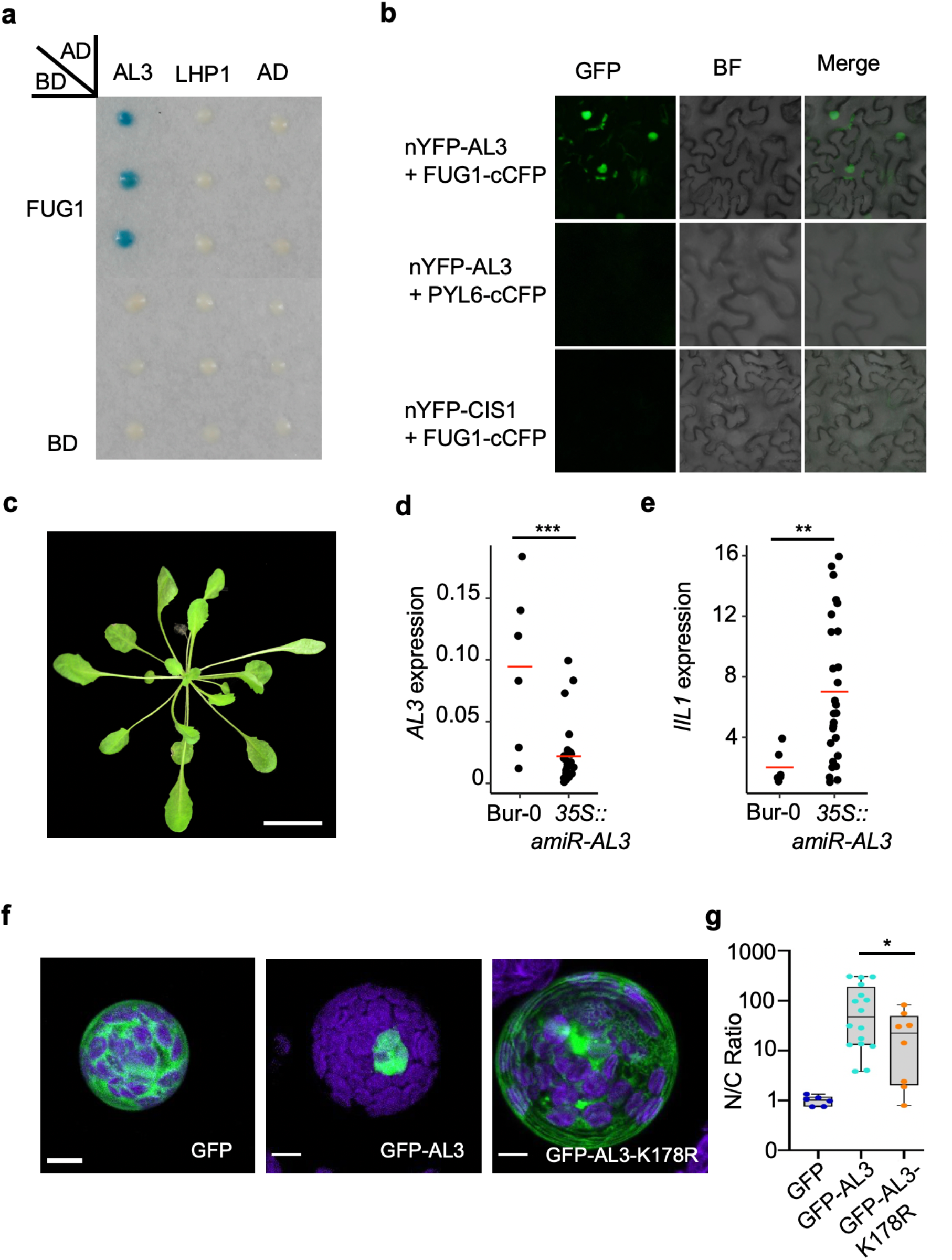
SUMO protease FUG1 interacts with histone reader AL3, which is required for repeat expansion-induced downregulation of *IIL1*. a) FUG1 interacts with AL3 but not LHP1 in yeast cells. b) FUG1 interacts with AL3 *in planta* in BiFC assays using split-GFP in tobacco epidermal cells. PYL6 and CIS1 are used as negative controls in these assays respectively for C and N terminal fusion constructs. GFP-GFP channel, BF-Bright field. c-e) Knocking down AL3 suppresses the *iil* phenotype in *35S::amiR-AL3* plants (c) with downregulation of *AL3* expression (d) corresponding increase in *IIL1* expression (e). Each dot represents individual wild type Bur-0 or *35S::amiR-AL3* primary (T_1_) transgenic lines. f) Disrupting the potential SUMOylation site in AL3 perturbs its nuclear localisation. Representative maximum intensity projections showing the localisation of GFP or GFP-AL3 or AL3 harbouring K178R mutation (GFP-AL3-K178R) in Bur-0. Scale bar = 2.5 μm. g) Box and whisker plots of N/C (nuclear:cytoplasm) ratio of GFP (blue) or GFP-AL3 (cyan) or GFP-AL3-K178R (orange) in Bur-0. Each dot represents the quantification from an individual protoplast expressing the corresponding transgene. Statistical comparisons were done with a Student’s t-test. P-values: **<0.01, ***<0.001, ****<0.0001.

We then asked whether this FUG1-AL3 interaction is linked with the role of FUG1 in repeat expansion-associated gene silencing. To assess this, we generated knockdown lines *(35S::amiR-AL3)* in the Bur-0 background. More than 90% of the *35S::amiR-AL3* T1 transgenic lines (n=70) exhibited suppression of the *iil* phenotype coupled with a decrease *AL3* levels along with an increase in *IIL1* expression (Fig 3c-e). These results confirmed the requirement of AL3 for the downregulation of *IIL1* caused by expanded repeats.

Members of the Alfin-like family including AL3 are known to be nuclear localised SUMOylated proteins^26,27^. Consistent with this, using transient assays in tobacco cells, we observed AL3 to be potentially poly-SUMOylated (Fig S5). SUMOylation can have multiple effects on protein function which include promoting nuclear localisation and stabilisation^28-30^. We reasoned that perturbing the SUMOylation site in AL3 could allow us to assess its impact. SUMOylation site prediction tools GPS-SUMO^31^, JASSA^32^ and SUMO-PLOT (https://www.abcepta.com/sumoplot) identified K178 to be a major site of SUMOylation. We mutated Lysine to Arginine (K178R) and checked whether it would impact AL3 localization using Quantitative Fluorescence Lifetime Imaging Microscopy (FLIM) (Fig 3f-g, S6). We observed that K178R perturbed the nuclear localisation of AL3 (Fig 3f-g, S6), which indicated a potential role for SUMOylation in the nuclear localisation of AL3. However, we do not rule out other ways by which SUMOylation can affect AL3 function or other possible sites that might undergo SUMOylation.

AL3 is a known histone reader that binds to H3K4me3^33^ and Alfin-like family members are known to interact with components of the polycomb repressive complex to cause a chromatin state switch^34^. One of the components of the PRC1 complex is LIKE HETEROCHROMATIN 1 (LHP1), whose human orthologue HP1 is known to interact with SUMO proteases and this interaction is essential for epigenetic silencing^35,36^. Therefore, we considered whether FUG1 or AL3 could interact with LHP1, a component of the PRC1 complex. While we failed to detect any interaction between FUG1 and LHP1, we observed AL3-LHP1 interaction in yeast two-hybrid assays (Fig 3a & 4a). Split-GFP assays in tobacco cells confirmed an *in planta* AL3-LHP1 interaction (Fig 4b).

**Figure 4.**
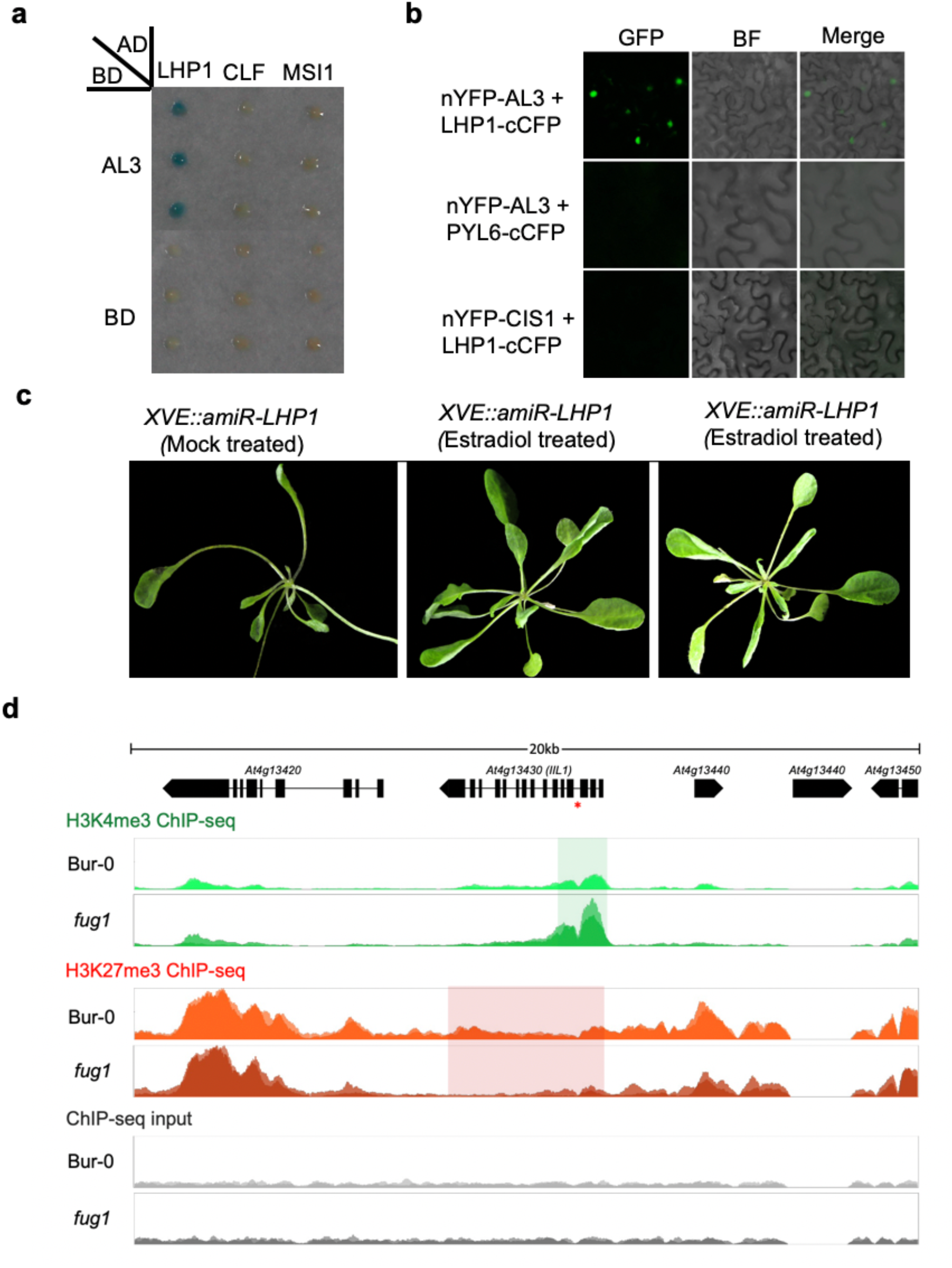
AL3 interacts with LHP1, a component of the PRC1 complex associated with the spread of H3K27me3 marks. a) AL3 interacts with LHP1 but not CLF or MSI in yeast cells b) AL3 interacts with LHP1 *in planta* in BiFC assays using split-GFP in tobacco epidermal cells. PYL6 and CIS1 are used as negative controls in these assays respectively for C and N terminal fusion constructs. GFP-GFP channel, BF-Bright field. c) Photographs of mock-treated or estradiol treated plants expressing artificial microRNAs against LHP1 (*35S::amiR-LHP1)* in the Bur-0 background. d) ChIP-seq profiles in a 20-Kb region surrounding *IIL1* for H3K4me3 or H3K27me3 marks compared with the input. Data from Coverage plots of reads across the *IIL1* locus (At4g13430, middle), normalised to the total number of mapped reads. The red asterisk in gene model shows the location of the GAA/TTC tandem repeat. Replicates are overlaid. H3K4me3 ChIP-seq coverage is shown in green. The green rectangle shows the boundaries of the peak called at *IIL1*, which shows significantly increased coverage in the *fug1* mutant. H3K27me3 ChIP-seq coverage are shown in orange. The red rectangle shows the gene body, which shows significantly decreased coverage in the *fug1* mutant. Input for the ChIP-seq samples are shown in grey.

We have previously tested whether knocking down LHP1 can suppress the *iil* phenotype^4^. Due to the strongly disrupted phenotype of *35S::amiR-LHP1*, we could not establish this link. To test this further, we generated estradiol-inducible knockdown lines of *LHP1* in the Bur-0 background *(XVE::amiR-LHP1)* and induced the artificial microRNAs against *LHP1* at the 7th leaf stage at 27ºC in short days. Of the 40 independent transgenic lines we observed a suppression of *iil* phenotype in 25 lines (Fig 4c) coupled with an increase in *IIL1* expression (Fig S7). These findings suggest that *LHP1* is required for epigenetic silencing caused by expanded repeats.

The interaction between FUG1-AL3/AL3-LHP1 raised the possibility that the epigenetic silencing might involve reading the H3K4me2/3 marks by AL3 followed by the recruitment of the PRC1 complex to cause the chromatin state switch to H3K27me3 and condense the chromatin. An earlier study has shown the *IIL1* locus to be a target for LHP1^37^. Therefore, we analysed the epigenetic status of the *IIL1* locus in the *fug1* mutant background by Chromatin Immunoprecipitation followed by sequencing (ChIP-Seq). We observed significantly high enrichment of H3K4me3 marks at the *IIL1* locus in *fug1* mutants compared to Bur-0 (Fig 4d). The enrichment was higher at the 5’ region of the gene including the transcriptional start site (TSS). Conversely, we also observed an increase in the H3K27me3 mark in Bur-0 compared to *fug1* mutants (Fig 4d). However, there is a spreading of the H3K27me3 throughout the *IIL1* locus, consistent with the involvement of LHP1. These findings indicate the requirement of FUG1 in modulating the epigenetic status of *IIL1* in presence of the repeat expansion, potentially involving a chromatin state switch by AL3/LHP1.

Our earlier findings indicated that triplet repeat expansions lead to 24-nt small RNAs, which in turn recruit the RdDM components to the *IIL1* locus resulting in epigenetic silencing. An initial transcription is required to trigger this response^4^. We have now demonstrated that in the final steps SUMO protease FUG1, histone reader AL3 and the PRC1 component LHP1 are required for epigenetic silencing caused by expanded repeats, which involves direct interactions between FUG1-AL3 and AL3-LHP1. We propose one possible model in which the FUG1 SUMO protease potentially deSUMOylates AL3. While it is unclear how deSUMOylation of AL3 affects its function AL3 interacts with LHP1 and recruits it to the *IIL1* locus thereby generating a chromatin state switch from H3K4me3 to H3K27me3 and allowing its spread resulting in gene silencing (Fig 5). Loss of any one of these components alleviates epigenetic silencing caused by expanded repeats (Fig 5). Consistent with this proposed model, we failed to detect small RNAs that map to the *IIL1* locus in the *fug1* mutant(Fig S8), similar to what we have previously reported in the knockdowns for *AGO4, DCL3* or *PolV*^4^. We however cannot rule out that FUG1 could target other RdDM components, since some of them are known to be SUMOylated^26^, though they were not recovered in our two-hybrid screen.

**Figure 5.**
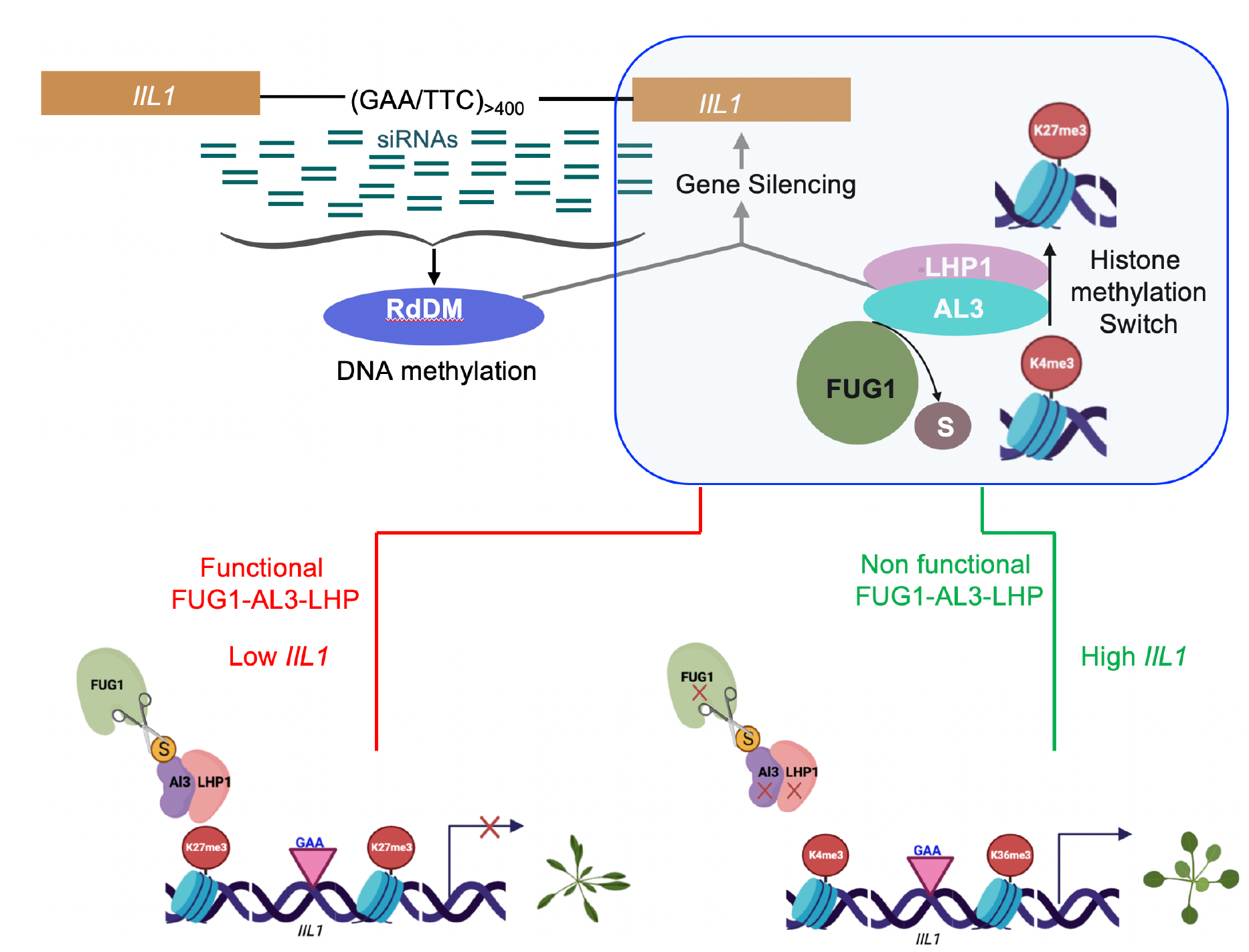
A model for epigenetic silencing caused by expanded repeats and the role of FUG1-AL3-LHP1 at the *IIL1* locus. Expanded repeats leads to small RNAs, which through the RdDM pathway targets the *IIL1* locus. This targeting involves changes in both DNA and histone methylations. The switch of histone H3K4me3 to H3K27me3 requires FUG1-AL3-LHP1 module. FUG1 interacts with AL3 and AL3 interacts with LHP1, which is essential for gene silencing. AL3-LHP1 interaction recruits LHP1 to cause H3K27me3 methylation. When FUG1-AL3-LHP module is functional, it results in repeat expansion-induced increase in H3K27me3 and epigenetic silencing and in the absence of AL3 or FUG1 or LHP1, this interaction with LHP1 does not occur and gene silencing is abolished, resulting in a normal plant.

In conclusion, we have demonstrated a key role for post-translational modifiers and histone readers in conferring epigenetic gene silencing in plants. Our studies draw parallels with the human system where SENP-mediated deSUMOylation of HP1 has been shown to be critical in its enrichment at pericentric chromatin^36^. Interestingly, in this case, while there is no direct interaction between the SUMO protease FUG1 and LHP1, a PHD finger protein AL3 acts as a conduit orchestrating a change in the chromatin state. While studies on differing species reveal some species-specific aspects, our study once again suggests that core principles of gene regulation may be conserved across eukaryotic systems. It will be interesting to see whether post-translational modifiers and histone readers have any role in the epigenetic silencing caused by expanded intronic repeats in Friedreich’s ataxia and other human diseases.

## Author contributions

Conceptualisation: SS and SB; Methodology: SS, HL, SA, AS, ER and SB; Software: CID, SS and SB; Formal analysis: SS, CB, JL, CID, CA, HMY, NS, GF, PT, PKB, ASY, BGB, SK, PS, OG, and SB; Investigation:, SS, JL, CA, HY, NS and CB; Writing – Original Draft: SS; Writing – Review and Editing: SS, AS, SA, HL and SB; Visualization: SS, SA, HMY and SB; Supervision: SS, RKY, SA, AS, HL and SB; Project administration: SS and SB; Funding acquisition: SS and SB.

## Acknowledgements

We thank Richard Clark (Utah, USA) for help with sequencing the *fug1* mutant, Andrej Seleznev for help with Bioinformatic analysis, Hiroaki Sako for initial characterisation of suppressors. We thank John Bowman, Jordyn Coutts, Yalong Guo, Rucha Sarwade, David Smyth and Lee Wong for discussions and comments on this manuscript. This work is supported by a Canadian Institute for Health Research grant PJT-178112 (ER), Department of Science and Technology – SERB Core grant (RKY), an NHMRC project grant APP1182090 (SB & SS), Australian Research Council Discovery Projects DP1095325 (SB) and DP190101818 (SB) and ARC Future Fellowships FT100100377 (SB), FT190100403 (SS), Australia-India Strategic Research Fund-Early and Mid-Career Fellowship (SS) and a Monash Larkins Fellowship (SB).

## Supplementary Figures

**Supplementary Figure S1.**
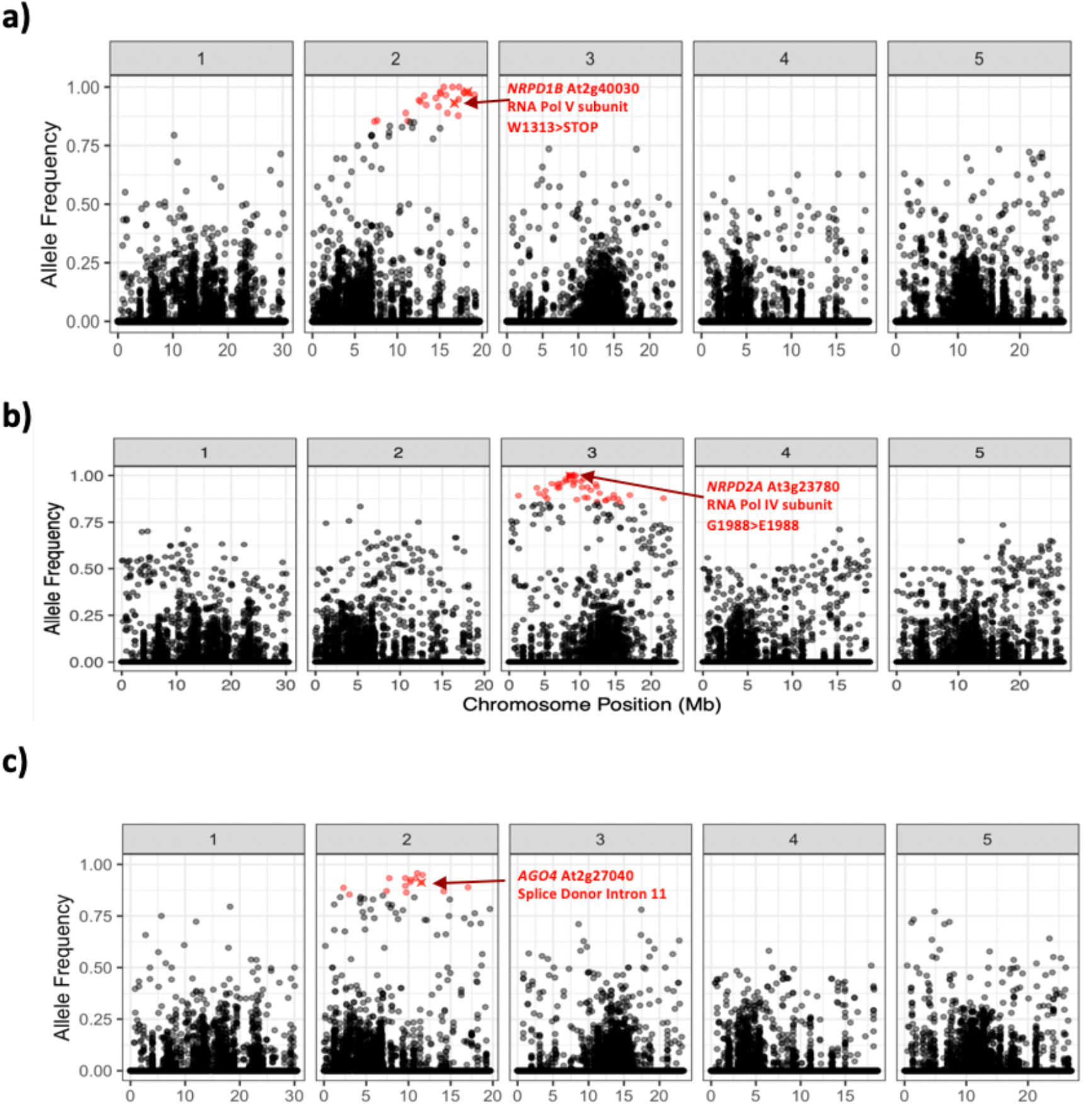
Genetic mapping of the suppressors of Bur-0 by SHOREMAP. Allele frequency of EMS-type SNPs in the fraction of F_2_ pooled plants that display the suppression of the *iil* phenotype. a) 49-9 b) 61-7 and c) 57-3. High frequency alleles (>0.85) are coloured red and red crosses show the putative causal alleles.

**Supplementary Figure S2.**
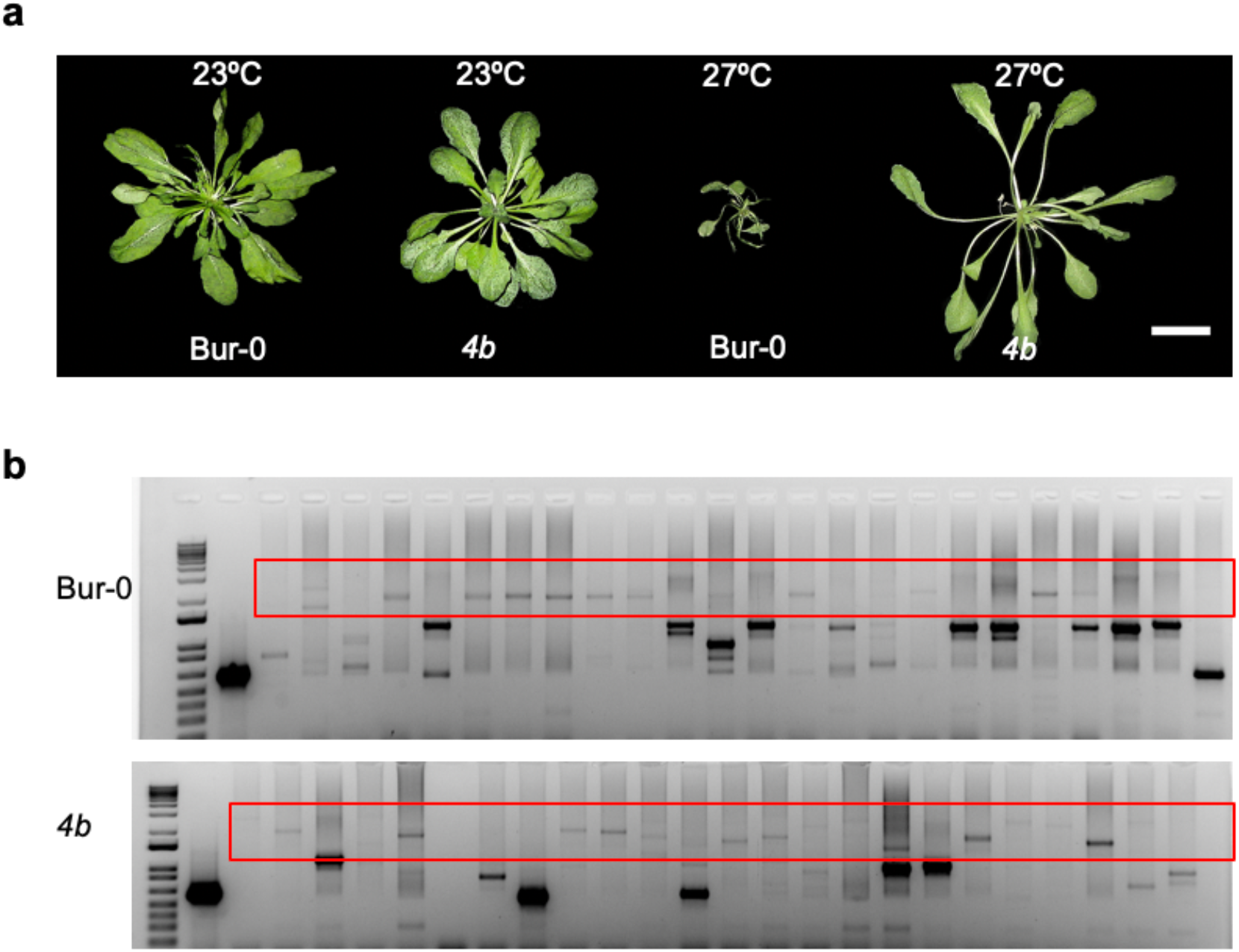
*fug1* mutant suppresses the temperature-dependent *iil* phenotype despite the presence of repeat expansion. a) *fug1* phenotype compared with Bur-0 at 23ºC and 27ºC. b) Suppression of the *iil* phenotype in *fug1* is not due to the loss of a repeat expansion. Gels showing retention of the expanded repeat in *fug1* plants compared with Bur-0 plants. The red boxes display the typical banding pattern of the repeat expansion. Each lane represents DNA analysed from individual plants with the Col-0 control in the first lane after the marker showing a non-expanded repeat.

**Supplementary Figure S3.**
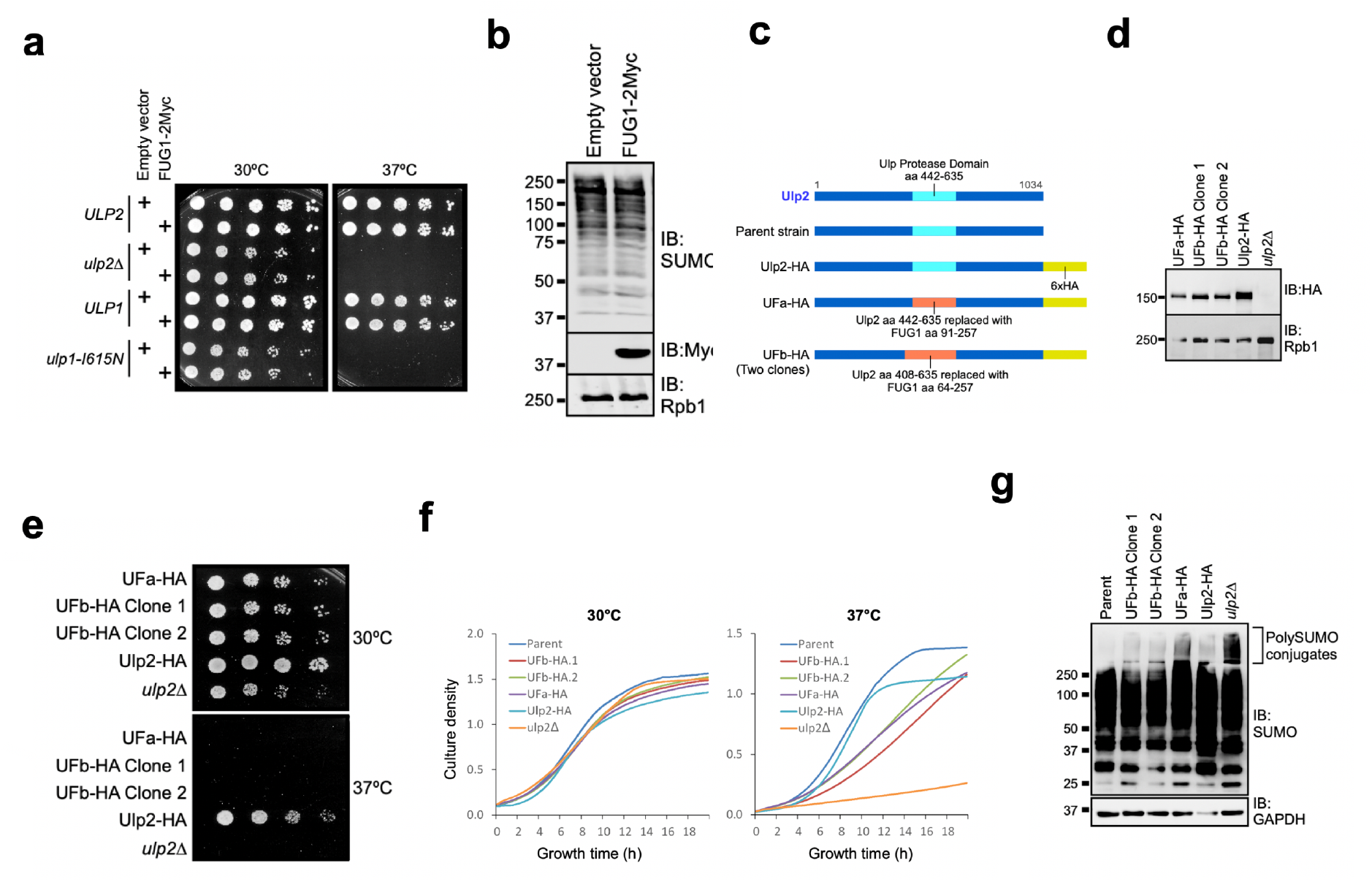
FUG1 is a functional deSUMOylase. a) Yeast growth assays in plates at 30ºC or 37ºC in wild type or mutants transformed with the empty vector of FUG1-2Myc constructs. FUG1 fails to rescue growth at 37ºC in the mutants. b) FUG1 is expressed, but does not show global alterations in the SUMO profile in yeast assays. c) Schematics of the two different domain swap experiments in which the protease domain of ULP2 is swapped with that of FUG1. d) Transgenes are expressed in yeast. e) Yeast growth assays in plates at 30ºC or 37ºC in wild type or mutants transformed with domain swapped hybrid constructs. Hybrid constructs do not rescue growth in plates. f) Partial rescue of the yeast mutants in cells transformed with the hybrid construct. g) Mutations in yeast ULP2 leads to accumulation of poly-SUMOylated proteins and the hybrid constructs reduce this poly-SUMOylated fraction indicating functional complementation.

**Supplementary Figure S4.**
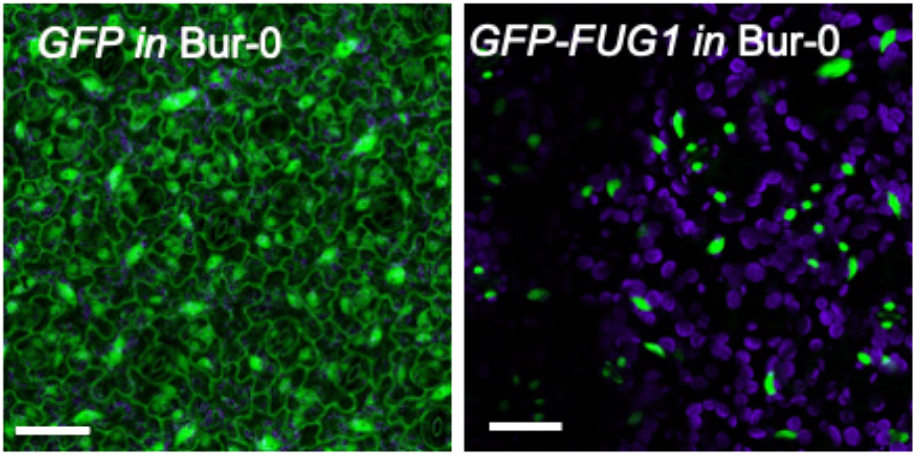
FUG1 is localised to the nucleus. Representative maximum intensity projections (MIPs) of protoplasts of transgenic Bur-0 leaves harbouring either GFP (left) or GFP-FUG1 (right). Autofluorescence (purple) was separated using fluorescence life time. Scale bar=10μM.

**Supplementary Figure S5.**
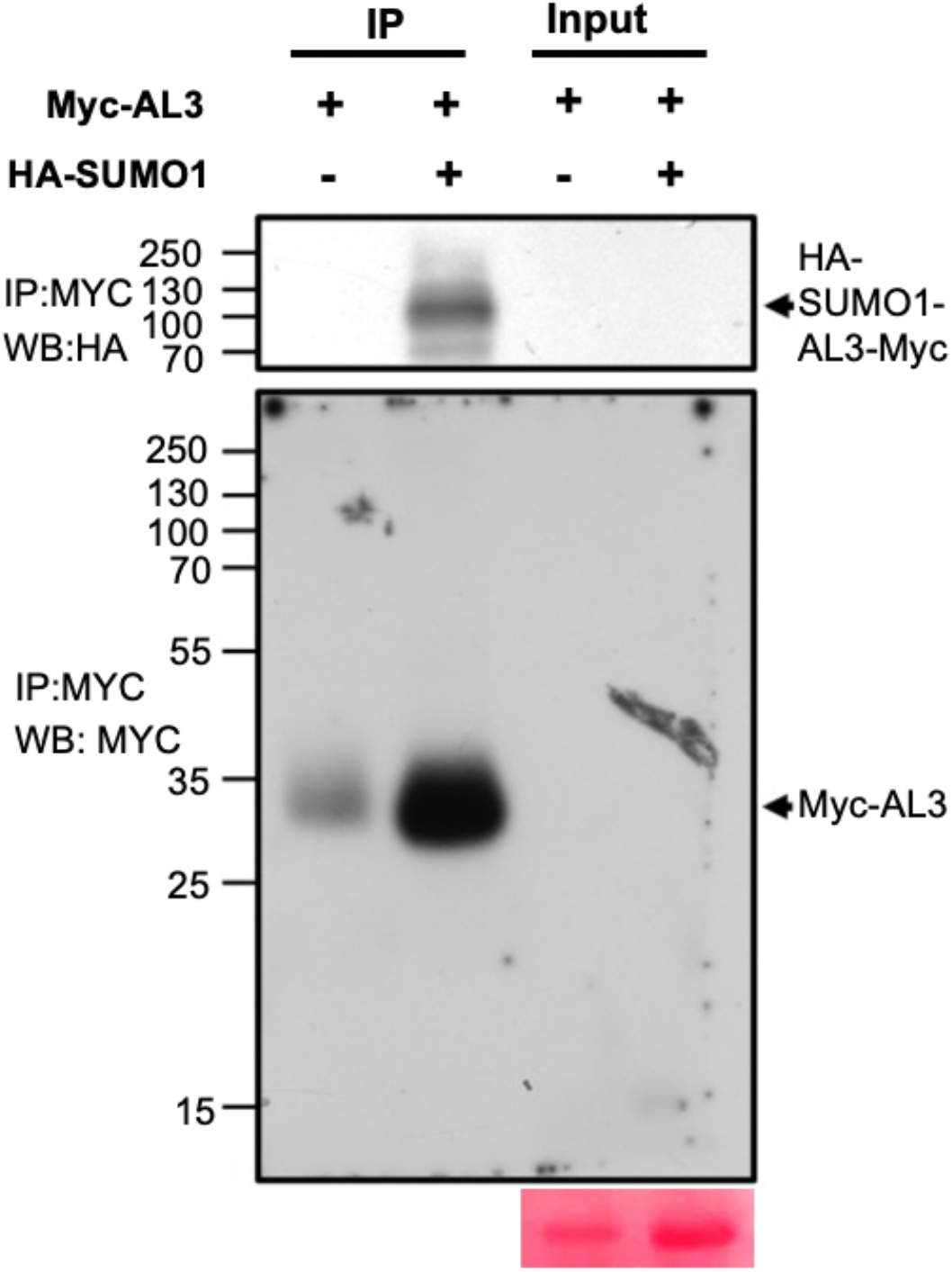
AL3 is SUMOylated in tobacco cells. SUMOylation assays with Myc-AL3 and HA-SUMO in tobacco cells. Blots were Immunoprecipitated with anti-Myc antibodies and blotted with either anti-HA antibodies (top) or anti-Myc antibodies (bottom). Ponceau S staining of input is shown for reference.

**Supplementary Figure S6.**
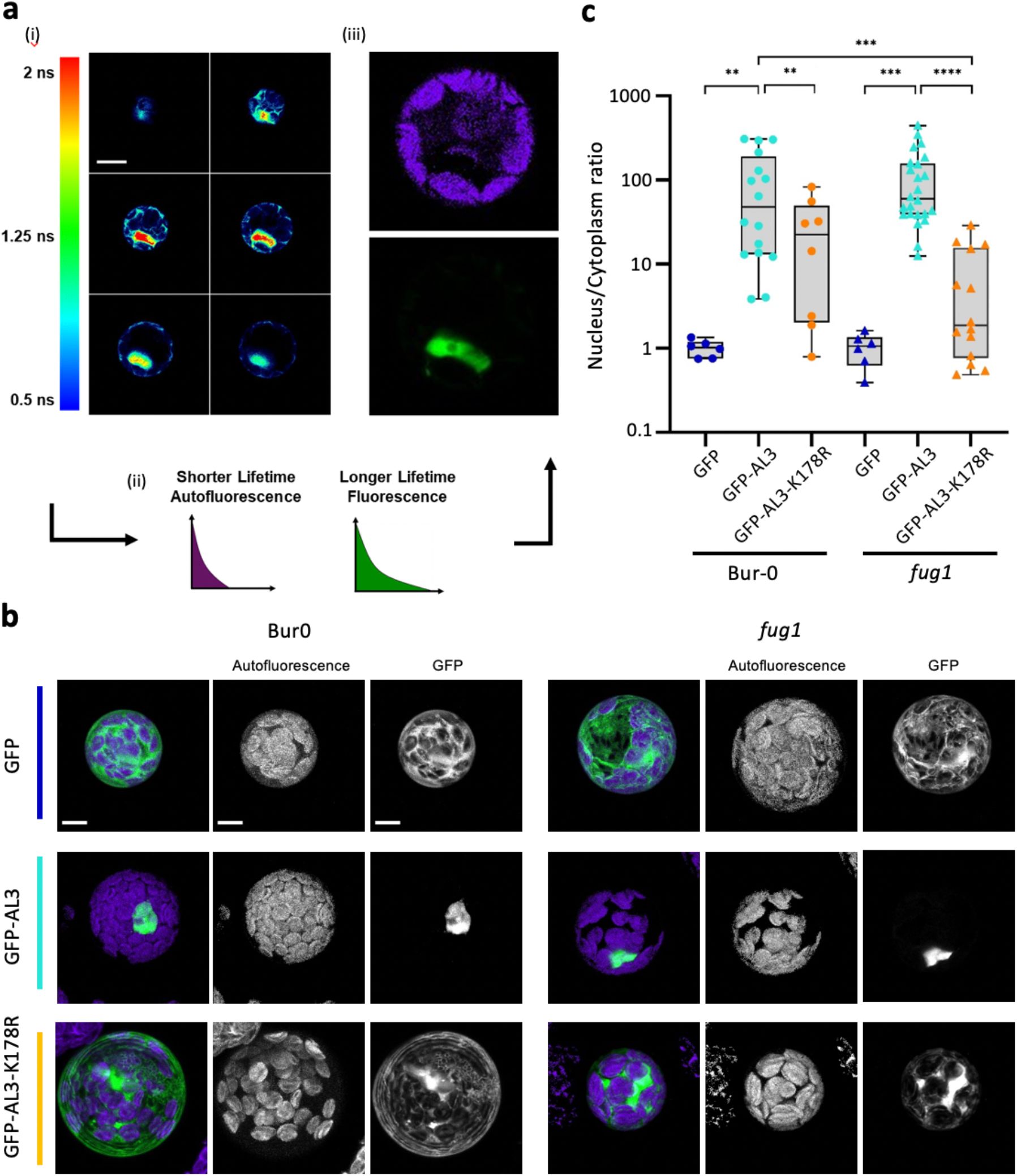
Perturbing the SUMOylation site of AL3 affects its sub-cellular localisation. a) A schematic of the Fluorescence Lifetime Imaging (FLIM) assay using transformed protoplasts isolated from Bur-0 or *fug1* mutants. Scale bar = 5 μm b) Representative maximum intensity projections showing the localisation of GFP or GFP-AL3 or AL3 harbouring K178R mutation (GFP-AL3-K178R) in Bur-0 and *fug1* mutant backgrounds 2.5 μm. c) Box and whisker plots of nuclear:cytoplasm ratio of GFP (blue) or GFP-AL3 (cyan) or GFP-AL3-K178R (orange) in Bur-0 and *fug1* mutant backgrounds. Each dot represents the quantification from an individual protoplast expressing the corresponding transgene. Statistical comparisons were done with a Student’s t-test. P-values: **<0.01, ***<0.001, ****<0.0001. The first lane of panel b and the left half (Bur) of panel c are also used in the Main Figure 3 (f &g).

**Supplementary Figure S7.**
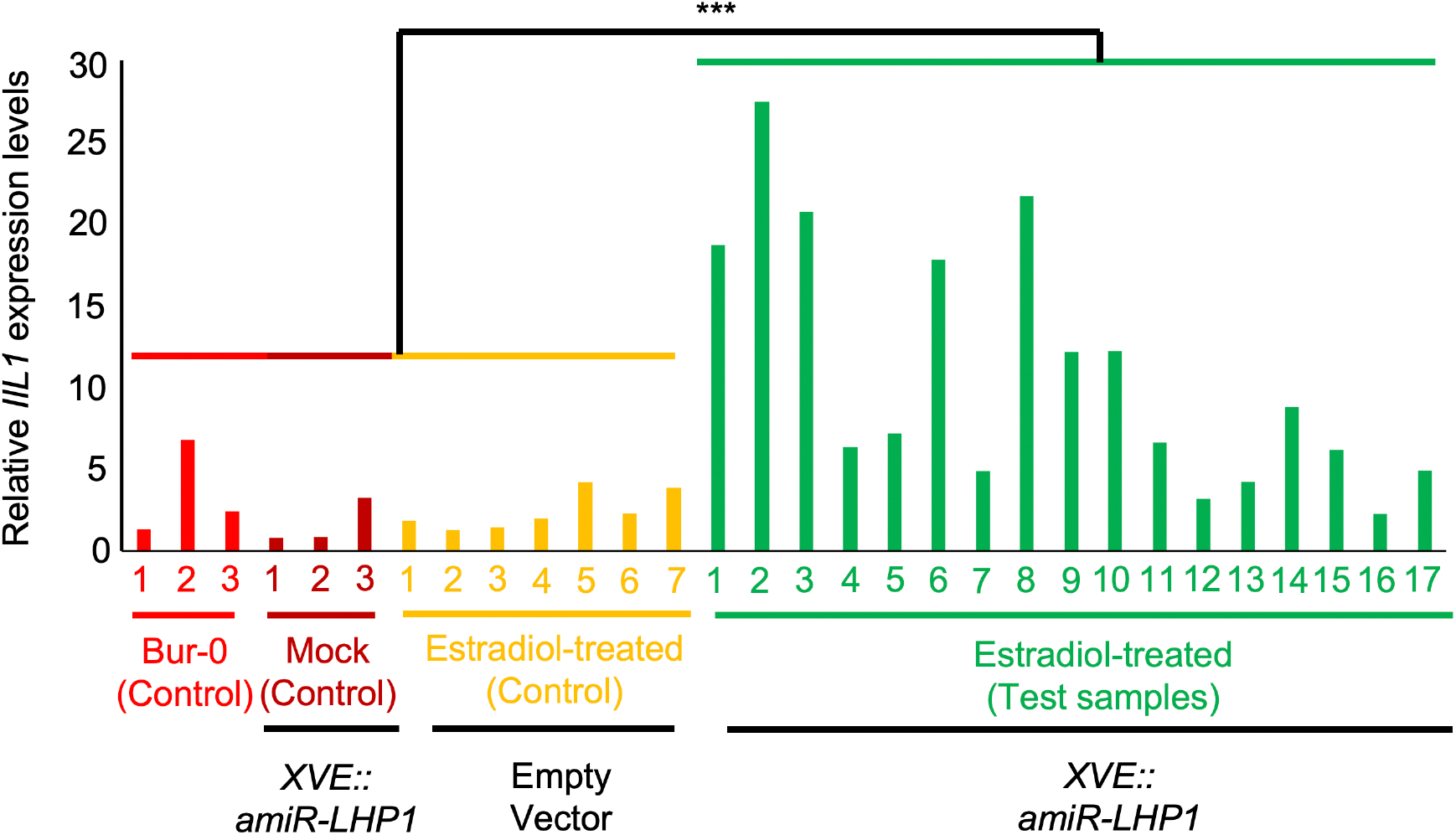
*XVE::amiR-LHP1* lines show an increase in *IIL1* levels upon estradiol induction. a) Relative *IIL1* expression levels. Comparison is made between Bur-0, empty vector controls as well as mock treated plants with that of estradiol treated samples in one-way ANOVA with the samples being separated as controls vs test. p-values: ***<0.0001.

**Supplementary Figure S8.**
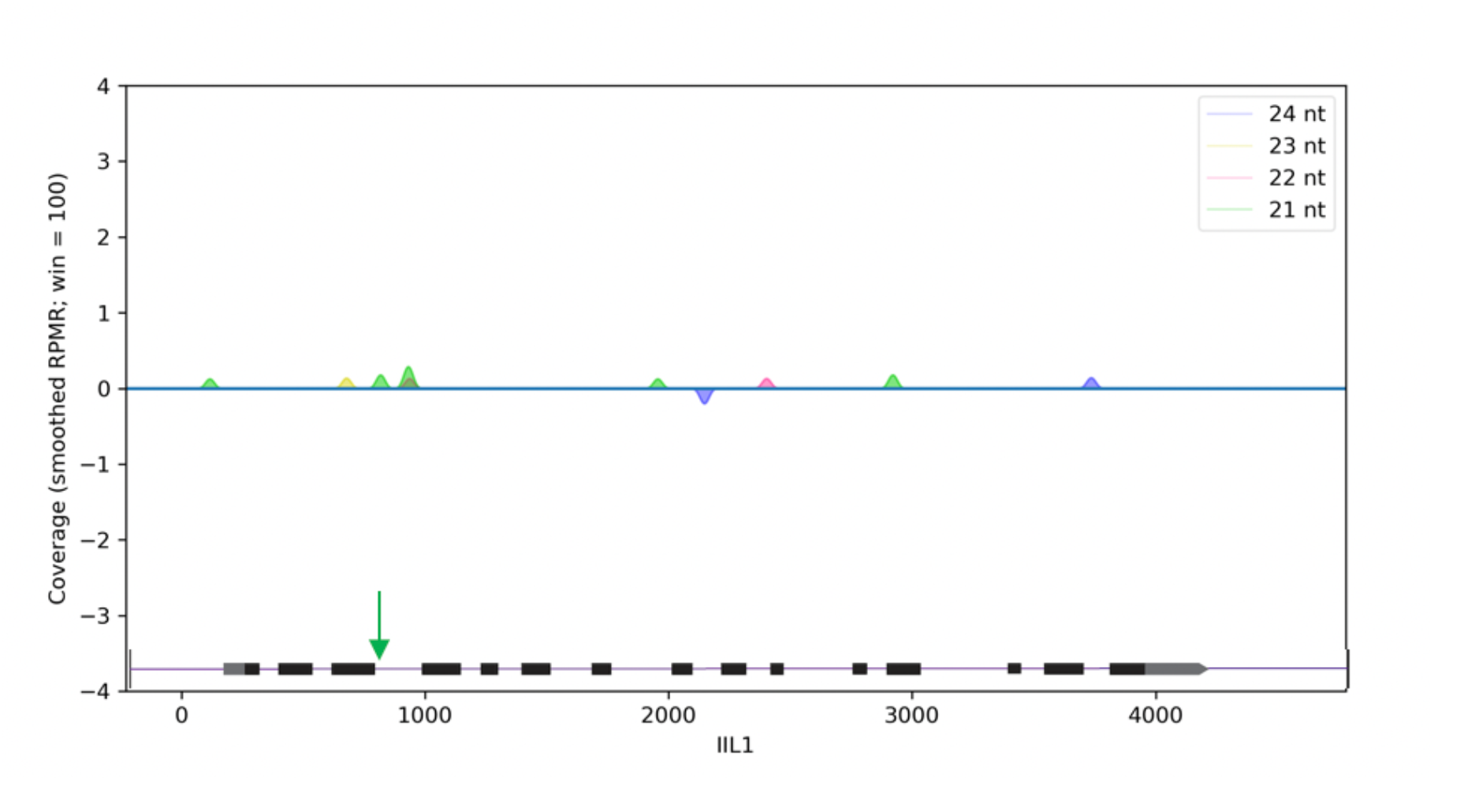
Abundance of siRNAs that map to *IIL1* locus in *fug1* mutant. Small RNAs that are typically found in Bur-0 ^4^ were not observed in the *fug 1* mutant. Small RNA profiles are generated as previously described^4^ using Small Complementary RnA Mapper (SCRAM) ^38^. The genic region is shown in the bottom with the black boxes and lines representing exons and introns, respectively. Normalised coverage along with standard error shown as shadows is shown for different types of small RNAs as indicated by the colour code. Only small RNAs that mapped to the non-triplet repeat sequences of *IIL1* are shown. Quantification of the sense and antisense small RNAs are shown in the positive and negative dimensions, respectively, along the y-axis. The green arrow indicates the position of the GAA/TTC repeat in the intron 3 of *IIL1*.

## Supplementary Tables

**Supplementary Table ST1.**
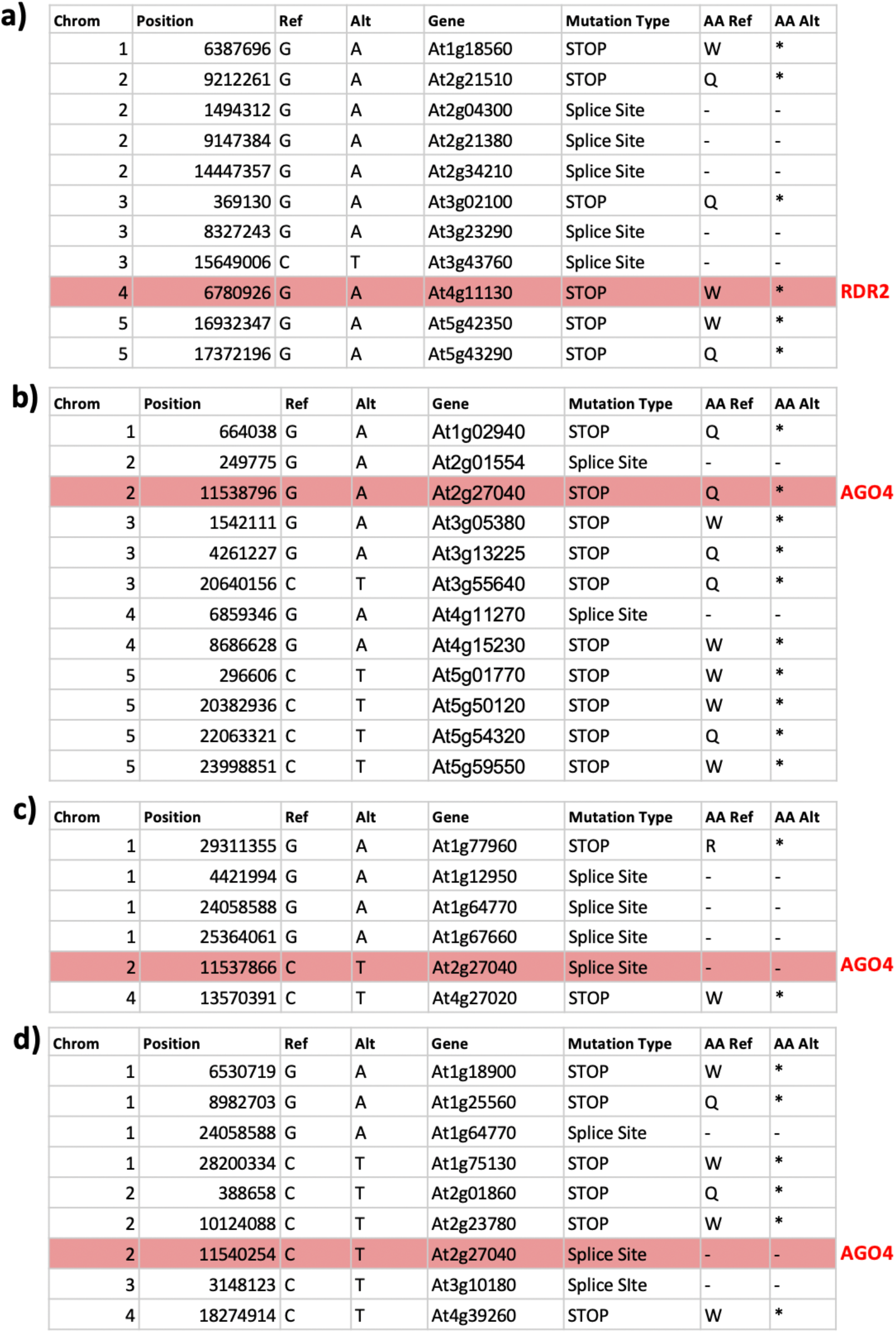
Candidate loss-of-function EMS-type SNPs in the phenotypic suppressors of Bur-0. a) 71-9 b) 76-2 c) 81-2 and d) 108-8. Mutations in the RdDM components are highlighted in red.

|  |  |  |  |  |  |  |  |  |  |
| --- | --- | --- | --- | --- | --- | --- | --- | --- | --- |
| **a)** | **Chrom** | **Position** | **Ref** | **Alt** | **Gene** | **Mutation Type** | **AA Ref** | **AA Alt** |  |
|  | 1 | 6387696 | G | A | At1g18560 | STOP | W | * |  |
|  | 2 | 9212261 | G | A | At2g21510 | STOP | Q | * |  |
|  | 2 | 1494312 | G | A | At2g04300 | Splice Site | - | - |  |
|  | 2 | 9147384 | G | A | At2g21380 | Splice Site | - | - |  |
|  | 2 | 14447357 | G | A | At2g34210 | Splice Site | - | - |  |
|  | 3 | 369130 | G | A | At3g02100 | STOP | Q | * |  |
|  | 3 | 8327243 | G | A | At3g23290 | Splice Site | - | - |  |
|  | 3 | 15649006 | C | T | At3g43760 | Splice Site | - | - |  |
|  | 4 | 6780926 | G | A | At4g11130 | STOP | W | * | **RDR2** |
|  | 5 | 16932347 | G | A | At5g42350 | STOP | W | * |  |
|  | 5 | 17372196 | G | A | At5g43290 | STOP | Q | * |  |
| **b)** | **Chrom** | **Position** | **Ref** | **Alt** | **Gene** | **Mutation Type** | **AA Ref** | **AA Alt** |  |
|  | 1 | 664038 | G | A | At1g02940 | STOP | Q | * |  |
|  | 2 | 249775 | G | A | At2g01554 | Splice Site | - | - |  |
|  | 2 | 11538796 | G | A | At2g27040 | STOP | Q | * | **AGO4** |
|  | 3 | 1542111 | G | A | At3g05380 | STOP | W | * |  |
|  | 3 | 4261227 | G | A | At3g13225 | STOP | Q | * |  |
|  | 3 | 20640156 | C | T | At3g55640 | STOP | Q | * |  |
|  | 4 | 6859346 | G | A | At4g11270 | Splice Site | - | - |  |
|  | 4 | 8686628 | G | A | At4g15230 | STOP | W | * |  |
|  | 5 | 296606 | C | T | At5g01770 | STOP | W | * |  |
|  | 5 | 20382936 | C | T | At5g50120 | STOP | W | * |  |
|  | 5 | 22063321 | C | T | At5g54320 | STOP | Q | * |  |
|  | 5 | 23998851 | C | T | At5g59550 | STOP | W | * |  |
| **c)** | **Chrom** | **Position** | **Ref** | **Alt** | **Gene** | **Mutation Type** | **AA Ref** | **AA Alt** |  |
|  | 1 | 29311355 | G | A | At1g77960 | STOP | R | * |  |
|  | 1 | 4421994 | G | A | At1g12950 | Splice Site | - | - |  |
|  | 1 | 24058588 | G | A | At1g64770 | Splice Site | - | - |  |
|  | 1 | 25364061 | G | A | At1g67660 | Splice Site | - | - |  |
|  | 2 | 11537866 | C | T | At2g27040 | Splice Site | - | - | **AGO4** |
|  | 4 | 13570391 | C | T | At4g27020 | STOP | W | * |  |
| **d)** | **Chrom** | **Position** | **Ref** | **Alt** | **Gene** | **Mutation Type** | **AA Ref** | **AA Alt** |  |
|  | 1 | 6530719 | G | A | At1g18900 | STOP | W | * |  |
|  | 1 | 8982703 | G | A | At1g25560 | STOP | Q | * |  |
|  | 1 | 24058588 | G | A | At1g64770 | Splice Site | - | - |  |
|  | 1 | 28200334 | C | T | At1g75130 | STOP | W | * |  |
|  | 2 | 388658 | C | T | At2g01860 | STOP | Q | * |  |
|  | 2 | 10124088 | C | T | At2g23780 | STOP | W | * |  |
|  | 2 | 11540254 | C | T | At2g27040 | Splice Site | - | - | **AGO4** |
|  | 3 | 3148123 | C | T | At3g10180 | Splice Site | - | - |  |
|  | 4 | 18274914 | C | T | At4g39260 | STOP | W | * |  |

**Supplementary Table ST2.** Major-effect mutations identified by sequencing *4b/fug1* mutant. The causal mutation in mapped interval is highlighted.

| Gene | Position | Ref | Alt | mRNA | Mutation type | Ref AA | Alt AA |
| --- | --- | --- | --- | --- | --- | --- | --- |
| Chr1 | 246854 | C | T | AT1G01680.1 | Splice-site |  |  |
| Chr1 | 4309533 | C | T | AT1G12660.1 | Nonsyn | Q | * |
| Chr1 | 5415300 | C | T | AT1G15750.1 | Splice-site |  |  |
| Chr1 | 5415300 | C | T | AT1G15750.2 | Splice-site |  |  |
| Chr1 | 5415300 | C | T | AT1G15750.3 | Splice-site |  |  |
| Chr1 | 5415300 | C | T | AT1G15750.4 | Splice-site |  |  |
| Chr1 | 11176753 | G | A | AT1G31260.1 | Nonsyn | Q | * |
| Chr1 | 18798131 | C | T | AT1G50730.1 | Nonsyn | Q | * |
| Chr1 | 26789566 | C | T | AT1G71015.1 | Nonsyn | W | * |
| Chr2 | 1068302 | G | A | AT2G03510.1 | Splice-site |  |  |
| Chr2 | 13136011 | G | A | AT2G30840.1 | Nonsyn | Q | * |
| Chr2 | 14983014 | G | A | AT2G35640.1 | Nonsyn | W | * |
| Chr2 | 16887625 | C | T | AT2G40435.1 | Nonsyn | Q | * |
| Chr3 | 162337 | G | A | AT3G01420.1 | Nonsyn | Q | * |
| Chr3 | 808948 | G | A | AT3G03400.1 | Nonsyn | Q | * |
| Chr3 | 5922801 | G | A | AT3G17340.1 | Nonsyn | Q | * |
| Chr3 | 5922801 | G | A | AT3G17340.2 | Nonsyn | Q | * |
| **Chr3** | **17958141** | **G** | **A** | **AT3G48480.1** | **Nonsyn** | **Q** | **\*** |
| Chr4 | 643756 | G | A | AT4G01510.1 | Splice-site |  |  |
| Chr4 | 7983675 | C | T | AT4G13750.1 | Nonsyn | Q | * |
| Chr4 | 7985383 | G | A | AT4G13750.1 | Splice-site |  |  |
| Chr4 | 9812602 | C | T | AT4G17610.1 | Nonsyn | W | * |
| Chr4 | 16303082 | C | T | AT4G34030.1 | Nonsyn | Q | * |
| Chr5 | 2051936 | C | T | AT5G06670.1 | Splice-site |  |  |
| Chr5 | 2431279 | C | T | AT5G07670.1 | Nonsyn | Q | * |
| Chr5 | 3603840 | C | T | AT5G11300.1 | Splice-site |  |  |
| Chr5 | 8327222 | G | A | AT5G24390.1 | Nonsyn | Q | * |
| Chr5 | 9566236 | C | T | AT5G27170.1 | Nonsyn | Q | * |
| Chr5 | 10825290 | G | A | AT5G28810.1 | Splice-site |  |  |
| Chr5 | 10925977 | G | A | AT5G28900.1 | Nonsyn | W | * |
| Chr5 | 17934666 | G | A | AT5G44520.1 | Nonsyn | Q | * |
| Chr5 | 19725625 | C | T | AT5G48640.1 | Nonsyn | W | * |
| Chr5 | 21164677 | G | A | AT5G52070.1 | Nonsyn | W | * |
| Chr5 | 22192936 | G | A | AT5G54630.1 | Nonsyn | R | * |

**Supplementary Table ST3.** Yeast strains used in this study.

| Description |  | Name | Genotype | Source |
| --- | --- | --- | --- | --- |
| ULP2 parent |  | BY4741 | *MATa his3Δ1 leu2Δ0 met15Δ0 ura3Δ0* | Open Biosystems |
|  | + empty vector | YER071 | + [pMGL3] | This study |
|  | + FUG1-2Myc | YER072 | + [pFUG1-2Myc] | This study |
| *ulp2Δ* |  | ERYM579 | *MATa his3Δ1 leu2Δ0 met15Δ0 ura3Δ0* | This study |
|  | + empty vector | YER073 | + [pMGL3] | This study |
|  | + FUG1-2Myc | YER074 | + [pFUG1-2Myc] | This study |
| ULP1 parent |  | MH1006 | *leu2-3,112 ura3-52 trp1-289 ade2Δ ade3Δ lys1::kanMX4* | Soustelle et al<sup>22</sup> |
|  | + empty vector | YER075 | + [pMGL3] | This study |
|  | + FUG1-2Myc | YER076 | + [pFUG1-2Myc] | This study |
| *ulp1-I615N* |  | MH1018 | *leu2-3,112 ura3-52 trp1-289 ade2Δ ade3Δ lys1::kanMX4 ulp1-I615N* | Soustelle et al<sup>22</sup> |
|  | + empty vector | YER077 | + [pMGL3] | This study |
|  | + FUG1-2Myc | YER078 | + [pFUG1-2Myc] | This study |
| Parent strain for hybrids |  | W303a | *MATa ura3-52 trp1Δ2 leu2-3_112 his3-11 ade2-1 can1-100* | Open Biosystems |
| UFa-HA |  | YBB053 | + *ULP2-[Δ442-635 + At FUG1 91-257]-6HA::Kl TRP1* | This study |
| UFb-HA |  | YBB054A,B | + *ULP2-[Δ408-635 + At FUG1 64-257]-6HA::Kl TRP1* | This study |
| Ulp2-HA (In YPH499 background) |  | ERYM513 | *ura3-52 lys2-801_amber ade2-101_ochre trp1-delta63 his3-delta200 leu2-delta1 ULP2-6HA::Kl TRP1* | This study |

**Supplementary Table ST4.**
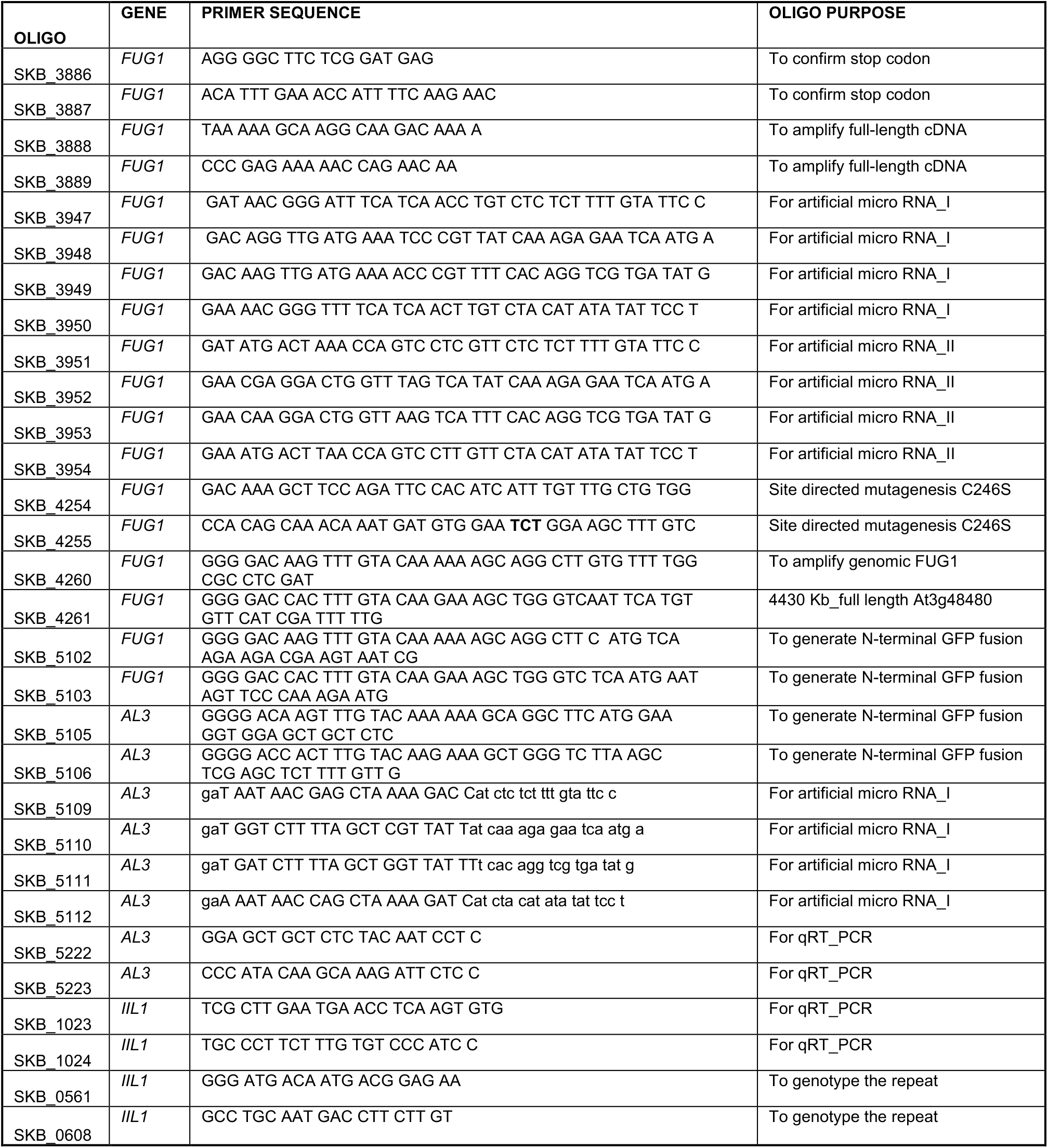
Primers used in this study.

## METHODS

### Plant material and growth conditions

*A. thaliana* accessions Bur-0 (and Pf-0 have been previously described ^5^. Plants were grown at 23 ºC and 27 ºC under short day (SD, 8 hours light, 16 hours darkness) conditions for DNA sequencing, CHIP-seq and Small RNA profiling. For the genetic suppressor screens approximately 32,000 Bur-0 seeds were mutagenized with ethyl methane sulfonate (EMS) as described previously ^5^. Approximately we pooled 10 M1 plants per family and the plants were grown at the Monash greenhouse in long day conditions for seed collection. In total we screened 120 M2 families at 27°SD to identify potential suppressors of the *iil* phenotype.

### Yeast two-hybrid screening library

Yeast two-hybrid screening was done as described previously^39^. The CDS of *FUG1* was cloned into the *pBridge* vector (Clonetech) and *FUG-pBridge* was used as the bait for Yeast two-hybrid screening. *FUG1-pBridge* and the plasmid library containing *pACT2* (from Dr. Joe Ecker, Salk Institute, California) were co-transformed into Y190 yeast strain for yeast two-hybrid screening. Diluted samples of the Y190 yeast strain were spread on appropriate SD selection agar plates (SD-Trp/-His/-Leu containing 30 mM 3-AT) and grown at 28ºC for 6-8 days. The mono-yeast clones with large volume were selected for strewing and X-Gal chromogenic reaction. The clones with blue X-Gal chromogenic reaction were used for yeast PCR, and the products of PCR were sequenced by sanger sequencing and NCBI Blast was used to identify the target genes. The CDS of *AL3* was cloned into the *pBridge* and *PGADT7* vectors (Clonetech), and the CDS of *LHP1, CLF, MSI1* was cloned into the *pBridge* and *PGADT7* vector, respectively. The different combinations were co-transformed into yeast AH109 or Y190 and plated on SD-Leu / -Trp medium for growth. Then the AH109 strains were transferred to SD-/-Trp/-His/-Leu/-Ala plate and the Y190 strains were subjected to X-Gal chromogenic reaction to test interaction.

### BiFC Assay

BiFC assays have been done according to Walter et al as described previously^39,40^. The CDS of *AL3* was cloned into the *pUC-SPYNE (nYFP-X)* vector. *FUG1* and *LHP1* were cloned into *pUC-SPYCE (cYFP-X)* vector. Each vector was transformed into Agrobacterium GV3101 and isolated single clones were incubated in LB at 28ºC for overnight. MES mixing buffer (10mM MgCl2, 10mM MES, 100μM acetobutylone) was used to dilute to suitable OD 600 (0.8∼1.0). The different combinations of GV3101 containing nYFP and cYFP were mixed respectively and infected the abaxial side of three-week-old tobacco leaves, and the fluorescence signals were observed by confocal 3 days later using an Olympus Fluorescence Microscope (Olympus CellSens Standard: Excitation wavelength 488nm; Emission wavelength 505nm).

### Subcellular localization analysis by Fluorescence lifetime imaging (FLIM)

Protoplasts and homozygous transgenic plants containing either EGFP, EGFP-AL3, EGFP-FUG1 or EGFP-AL3^K178R^ were imaged using an SP8 Falcon (Leica Microsystems) with an 86x 1.2 NA objective. Protoplasts were extracted using the whole leaf tape sandwich method as per Wu et al^41^ and transfections were carried out according to Yoo et al^42^. The protoplasts and leaves were imaged as volumes with slices taken at 0.5 μm and 2 μm respectively using a 1.0 AU pinhole. Images were acquired with 488 nm excitation at 10% transmission using a tuneable pulsed white-light laser. Emissions were collected from 500-560 nm using a HyD detector. The GFP and auto-fluorescence were separated using pixel-wise two-component fitting with τ1 = 2.600 ns and τ1 ≈ 0.2 ns. These images were analysed as volumes and the representative images were displayed as maximum intensity projections.

### SHOREMAP analysis and sequencing of mutants

For bulk segregent analysis we pooled leaf tissues from more than 500 plants that displayed the suppressor phenotype (normal-looking) from the segregating F_2_ population derived from a backcross with the Bur-0 parent. Other mutants were directly sequenced. Genomic DNA was extracted by using DNeasy Plant Maxi kit (cat.no.68163) from Qiagen. All Quality controlled (QC) passed samples were processed for sequencing on the Illumina Novaseq PE 150 platform by GENEWIZ-China. Approximately 30million reads/sample were analysed.

Quality-controlled sequences were aligned to the TAIR10 reference genome using Bowtie2 (v2.4.4)^43^. Called SNPs using samtools (v1.13; mpileup -E -uf) and generated VCF files with bcftools (v1.13; call -c). We then ran SHOREMap (v3.6) to identify SNPs most enriched in suppressor-phenotype F_2_ plants (foreground) compared to the Bur-0 parent (background). We found variant SNPs in suppressor-phenotype F_2_ VCFs using SHOREmap (v3.6; convert command). We were unable to run the SHOREmap extract command, so we developed a python script (filter_subtract_variants_vcf.py) to calculate the allele frequencies as follows: Allele Frequency = Frequency of SNP in foreground -Frequency of SNP in background. E only considered SNPs if they were EMS-type C>T or G>A mutations. SNPs were considered putatively loss-of-function if they generated a stop codon, or altered the core sequence (GT or AG) of an annotated splice site; or otherwise if one of the most enriched SNPs was predicted to cause a change in amino acid sequence.

Given that our first four suppressors mapped to genes in the RdDM pathway, we adapted our pipeline to identify candidate genes without requiring a backcross. For the suppressor mutants themselves: DNA was extracted, sequenced, aligned and processed to VCF files as above. We then ran SHOREMap (v3.6) to identify SNPs most enriched in the suppressors themselves (foreground) compared to the Bur-0 parent (background). This meant that all homozygous EMS SNPs unique to the suppressor were assigned an allele frequency of 1.0. We then took only those SNPs putatively causing a loss-of-function. We then asked if any of the dozen or so remaining SNPs belonged to known RdDM pathway genes.

### Small RNA sequencing

Small RNA sequencing has been previously described^4^. Leaf tissue from 45-day old *fug1* mutant plants were collected 5 hours after the beginning of the day (light exposure) and snap frozen immediately using liquid nitrogen. Total RNA was extracted using TRIzol reagent (Ambion). Quality controlled reads were mapped to the TAIR10 *Arabidopsis* genome using SCRAM^38^ and analysed as described previously^4^.

### ChIP-seq

For ChIP-seq Bur-0 and *fug1* mutant plants were grown for five weeks at 27°SD. 1.5 grams of leaf tissues from two independent biological replicates were used for chromatin preparation. Further chromatin preparation steps were followed as previously described^44^. We assessed two histone marks H3K4me3 and H3K27me3 and the ChIP grade antibodies against for these marks procured from Diagenode (H3K4me3, C15410003-50, and H3K27me3,C15410195). Purified DNA quality was checked by Bioanalyzer and QC passed samples were subjected to library preparation and proceeded for sequencing using Illumina Novaseq PE150 sequencing platform. Roughly 20M/6G per sample raw data was used for analysis. Clean reads were aligned to the TAIR10 genome using Bowtie2^43^ with default parameters. For H3K4Me3 samples, peaks were called using MACS3^45^ v3.0.0a7; Parameters: -f BAMPE, --broad, --broad-cutoff 0.1, -g 1.35e8). We took all peaks observed across all samples; where two peaks in different replicates overlapped, we considered them to be the same peak and took the maximal boundaries. For H3K27Me3 samples, we simply measured within TAIR10 annotated gene boundaries. Reads across regions (either peaks or genes) were counted using featureCounts from the Subread package^46^. Differentially covered regions were identified using DESeq2^47^.

### Yeast SUMOylation assays

Yeast strains used in this study are listed in Supplementary Table ST3. The FUG1-2Myc yeast expression plasmid was generated by PCR amplifying the *FUG1* cDNA with a 3′ 2x Myc-encoding sequence and sub-cloning into the BamHI restriction site of vector pMGL3. Ulp2-FUG1 hybrid-expressing strains were generated by fusion PCR of a DNA fragment that encodes residues 91 to 257 or 64 to 257 of *FUG1* and a fragment encoding a downstream portion of *Ulp2* fused to a 6xHA-encoding sequence and a *K. lactis TRP1* marker gene. The plasmid and PCR products were then transformed into appropriate yeast strains as previously described^48^. Yeast growth spot assays, preparation of extracts, and immunoprecipitation analysis were performed as previously described^49^.

### Transient In-vivo SUMOylation assay of AL3

To analyse the SUMOylation of AL3, PCR_ amplified full length cDNA of AL3 was cloned into pGWB621.The HA-SUMO construct has been previously described ^50^. Agrobacterium harboring AL3-myc and HA-SUMO1 were diluted in 10mM MgCl2 supplemented with 150μM acetosyringone with a final OD600 to 0.2 and co-infiltrated in 4 weeks old tobacco leaves as described previously^51^. After 3 days post infiltration, approximately 2g of the leaf samples were harvested and frozen in liquid nitrogen and stored in –80 until use. Total protein was extracted in SUMO extraction buffer (50mM Tris pH 8.5, 150mM NaCl, 1mM EDTA, 0.1% SDS, 1% NP-40, 20mM NEM, 0.5% Sodium deoxycholate and 1X Protease inhibitor cocktail) in cold room. The supernatant was incubated with anti-Myc magnetic beads for 30 minutes in a rotary mixer in a cold room. The bound protein was washed twice with extraction buffer and eluted in 1X samples loading buffer pre-heated at 95°C. Eluted protein was further analyzed in 10% SDS-PAGE and further transferred on polyvinylidene difluoride (PVDF) membrane. After blocking with 5% skimmed milk, membrane was further incubated with anti-HA (Abcam)/anti-SUMO1 (in-house produced) and anti-MYC (Sigma) overnight. After incubating the membrane with secondary antibodies for 1 hours, the membrane was vigorously washed and further developed the X-ray films by automated developer machine.

### Artificial microRNAs

Online tool WMD3 (http://wmd3.weigelworld.org/cgi-bin/webapp.cgi) was used to designing artificial miRNAs against *FUG1, Al3* and *LHP1. FUG1* and *LHP1* amiRNAs were generated by site directed mutagenesis as per Schwab et al^52^ and the *LHP1* amiRNA construction details were explained previously^4^. *AL3* amiRNA constructs were commercially synthesized in plasmids (IDT, USA) and then sub-cloned into Gateway compatible entry vector (pDONR221). The final artificial miRNA precursors were sub cloned into the plasmid pFK210 by conducting LR reactions according to the manufacturer’s protocol (Gateway® LR Clonase® II Enzyme mix, Invitrogen). The *LHP1* artificial microRNA was cloned into the Gateway-compatible pGREAT XVE-system vector^4^. Sequence verified constructs were transformed into *A. tumefaciens* GV3103 further transformed into Col-0, Bur-0, and *fug1* genotypes using the floral dip method ^53^. The first generation (T_1_) of transformants were grown at 27 ºC SD and continuously watered with nutrient water containing 120 mg/L BASTA (glufosinate ammonium, BAYER) for selection. For the inducible system, plants were sprayed with 50μM Estradiol with 0.1% Silwet L-77, every two days after the emergence of the 7^th^ leaf, as previously described^4^. We obtained multiple independent primary transgenic lines and most of the transgenic lines displayed phenotypic suppression. The number of primary independent transformants that displayed complete phenotypic suppression for each of the transgenes *35S::amiR-FUG1 –190,35S::amiR-AL3 –68);; 35S::XVE::amiR-LHP1-25)*. The presence of each transgene was confirmed by PCR using the primers listed in Supplementary Table ST4 and the expression level of each artificial miRNA’s target gene was determined by qRT-PCR. Expanded GAA/TTC repeat tracts at the *IIL1* locus were confirmed by PCR (primer pair: oSKB_608 and oSKB_561, Supplementary Table ST4). After phenotypic analysis at 27 ºC short days, primary transformants were transferred to 23 ºC long day conditions for seed collection.

### Expression analysis

For qRT-PCR, total RNA was extracted using TRIzol reagent (Ambion). cDNA was synthesized with Anchored-oligo (dT)_18_ primers and the Transcriptor First Strand cDNA Synthesis Kit (Roche). cDNA was diluted five-fold with water and 2-4μl of the diluted cDNA served as template for each qRT-PCR as described previously ^4^. Variation in gene expression was analyzed through quantitative real-time PCR analysis using the 2^-ΔΔcT^ method^54^. The statistical significance of the difference in gene expression between specific samples is analyzed through Student’s *t*-test or ANOVA. For ChIP experiments, the data is expressed after normalizing with a positive control and the statistical significance was analyzed through a Student’s *t*-test.

## ADDITIONAL RESOURCES

All sequencing data are available at NCBI short reads archive. The accession number for the ChIP-seq reads is XXXXX and the accession number for the small RNA sequence from *fug1* is XXXXX.

